# Cell-free Protein Crystallization for Nanocrystal Structure Determination

**DOI:** 10.1101/2022.04.15.488232

**Authors:** Satoshi Abe, Junko Tanaka, Mariko Kojima, Shuji Kanamaru, Kunio Hirata, Keitaro Yamashita, Ayako Kobayashi, Takafumi Ueno

## Abstract

In-cell protein crystallization (ICPC) has attracted attention as a next-generation structural biology tool because it does not require multistep purification processes and large-scale crystallization screenings. However, significant issues remain to be solved in context of obtaining various protein crystals in sufficient amounts and quality for structure determination by ICPC. Here, we report the development of cell-free protein crystallization (CFPC), a direct protein crystallization technique which uses cell-free protein synthesis. The most crucial advantages of CFPC are that the reaction scale and time can be minimized and that various reagents can be added during the reaction. We obtained high-quality nano-sized polyhedra crystals, which are produced in insect cells by infection with cytoplasmic polyhedrosis virus, at a 200 μL reaction scale within 6 h. We applied this technology to structure determination of crystalline inclusion protein A (CipA) by suppressing twin crystal formation with addition of an inhibitor to the reaction solution. We succeeded in determining a 2.11 Å resolution structure from the nanocrystals of CipA. This technology, which integrates in-cell and *in vitro* crystallizations significantly expands the tools available for high throughput protein structure determination, particularly in context of unstable, low-yield, or substrate-binding proteins, which are difficult to analyze by conventional methods.

## Introduction

Proteins crystallized in living cells have been frequently reported over the last few decades^1–3^. Such crystals provide biological functions such as protein storage, protection, heterogeneous catalysis, and activation of the immune system^4^. The relationships between the functions and structures of the crystals have been investigated by direct structure determination of the micron-sized crystals grown in living cells since the structure of polyhedra, a natural in-cell crystal, was first determined in 2007^5^. Thus, other than material applications of natural in-cell protein crystals, in-cell crystallization of various proteins has been widely expected to be developed as a next-generation structural biology tool because it does not require purification procedures and large-scale crystallization screening to obtain high-quality crystals^2^. In 2013, the crystal structure of cathepsin B from *Trypanosoma brucei* was determined using in-cell protein crystallization (ICPC) as the first example of determining the crystal structure of a recombinant protein^6^. Since then, ICPC has been attempted numerous times but structures of only a few proteins have been reported^4, 7–11^. This is because crystals are often incidentally formed in cells and their size and quality are insufficient for structural analysis^12^. Therefore, it appears that several technical issues must be overcome in applying this method for protein structure analysis.

Several ICPC methods, such as high throughput screening (HTS) and optimization of cell culture processes have been developed in efforts to resolve these problems^13–15^. A pipeline containing protein crystallization using insect cells with sorting by flow cytometry has been developed^15^. LaBaer et al. constructed a set of baculovirus expression vectors for a large-scale parallel expression of proteins in insect cells and successfully prepared microcrystals^14^. Mammalian and insect cells, currently used for ICPC, continue to represent a significant limitation as platforms to produce large numbers of high-quality microcrystals rapidly. Although another attempt has been made to add chemical reagents used for *in vitro* crystallization, it has not led to improvements in ICPC, possibly because the efficiency with which the reagents penetrate cell membranes and their effects on other cellular functions are unknown^12^. *Bacillus thuringiensis* (Bt) bacteria have been used to express cry protoxins as crystallization vessels for cargo proteins recombinantly, but the structures could not be determined^16, 17^. Once a new ICPC has been established and integrated with *in vitro* crystallization methods to overcome these concerns about ICPC, protein crystallography is expected to become a more accessible technology.

Cell-free protein synthesis (CFPS), a protein preparation technique used in synthetic biology, is very effective for rapid screening of protein synthesis^18^. However, it has been considered unsuitable for structural biology efforts that require large amounts of protein, such as crystallization^19–21^. Here, we report the application of CFPS to ICPC (Figure 1). We focus on (i) establishing crystallization of a protein using CFPS with small and rapid reactions and (ii) manipulating the crystallization by adding chemical reagents. The polyhedra crystal (PhC) produced in insect cells by infection of cypovirus (cytoplasmic polyhedrosis virus, CPV) is one of the most highly studied in-cell protein crystals. We obtained nano-sized PhCs in a 200 μL reaction within 6 h and succeeded in determining the protein structure at high resolution using a current standard beamline (BL32XU) at SPring-8, a large synchrotron facility.

**Figure 1.**
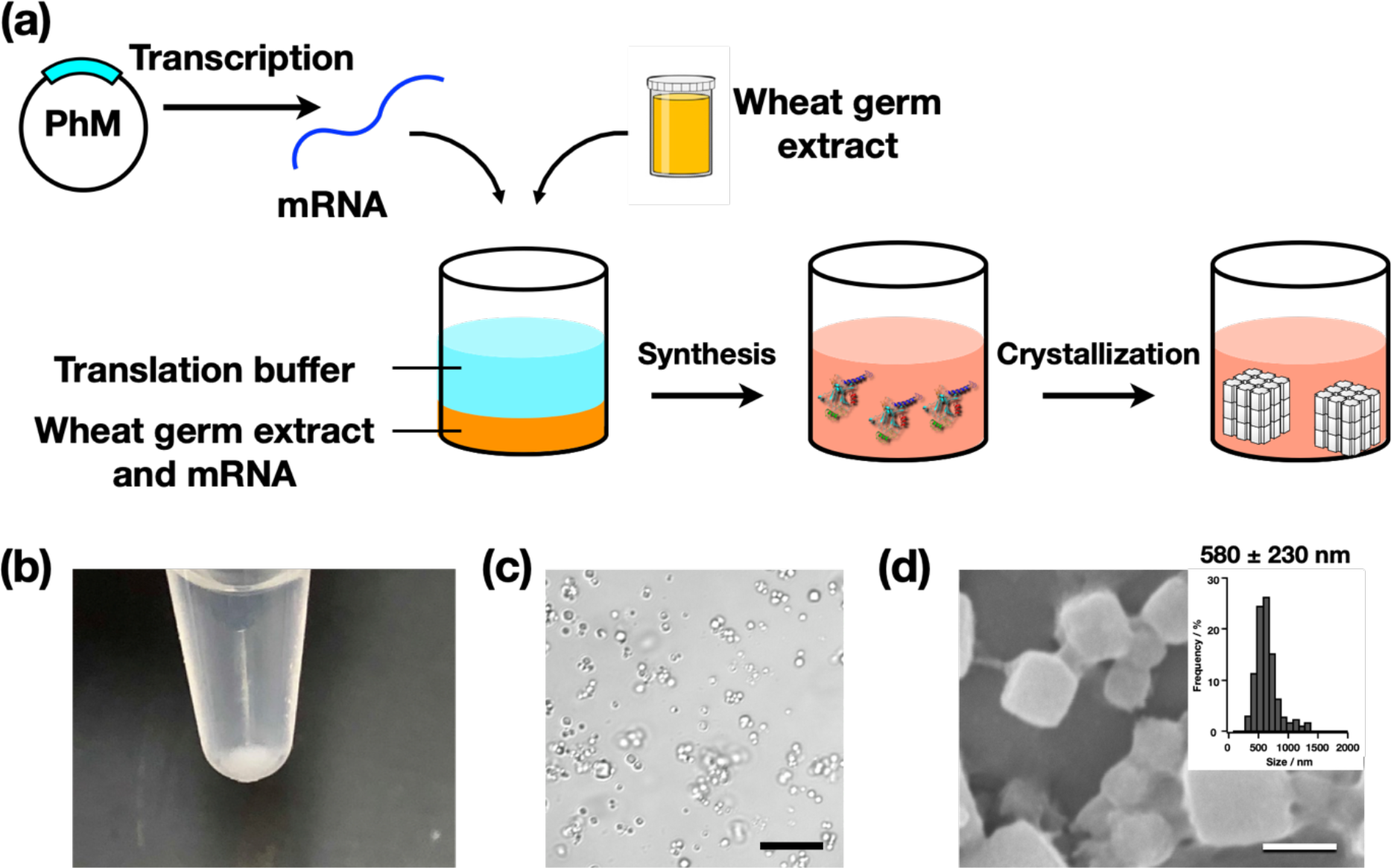
(a) Schematic illustration of Cell-Free Protein Crystallization (CFPC) of polyhedrin monomer (PhM) using the Wheat Germ Protein Synthesis kit. (b) Photograph of the tube after CFPC. (c) Differential interference contrast (DIC) image of **PhC_CF**. Scale bar = 10 μm. (d) A scanning electron micrograph of **PhC_CF**. Size distribution of **PhC_CF** determined by the SEM image. Scale bar = 1 μm.

Moreover, the most crucial advantage of CFPC is that various reagents can be added to the reaction mixture without preventing protein synthesis. The structure of crystalline inclusion protein A (CipA), a bacterial in-cell crystal, had not been previously reported^22^. Since we found that twin crystals are formed when CipA is expressed in *E.coli*, we applied CFPC to CipA with the addition of twin crystal inhibitors during the crystallization process and succeeded in obtaining suitable crystals and determining the structure of CipA at a 2.11 Å resolution. Therefore, CFPC, a hybrid method of ICPC and *in vitro* protein crystallization, can be developed at a surprisingly small scale to provide rapid crystallization without any purification procedures. CFPC opens up a new method for crystallizing unstable proteins and rapidly determining their structures.

## Results

### Crystallization of PhC by CFPC method

CFPS is a conventional synthetic biology method for protein structural determination^21^. It enables synthesis of proteins, such as membrane proteins and protein assemblies, that are difficult to purify using living cells^23, 24^. We performed CFPS using extracts from wheat germ because these extracts have been identified as having the highest protein expression activity among the eukaryotic systems^25^. Crystallization of polyhedrin monomer (PhM) was performed using the Wheat Germ Protein Synthesis kit (WEPRO^®^7240 Expression Kit). The translation reaction was carried out using the bilayer method. A 20 μL reaction mixture containing 10 μL of WEPRO^®^7240 and 10 μL of the mRNA solution was overlaid with a 200 μL SUB-AMIX^®^ SGC solution in a 1.5 mL microtube and then incubated at 20 °C for 24 h. White precipitates were collected after centrifuging the reaction solution (Figure 1b). The crystalline precipitate was observed with an optical microscope (Figure 1c). SDS-PAGE and matrix assisted laser desorption ionization-time of flight mass spectrometry (MALDI-TOF MS) of the precipitate showed a band at 28 kDa and a peak of 28,361 Da, respectively (Supplementary Figure 1). These results are consistent with the calculated molecular weight of the PhM (28,368 Da). The crystals prepared from the CFPC reaction (**PhC_CF**) have the same cubic morphology as that of PhC synthesized in insect cells (PhC_IC). The average size of **PhC_CF** (580 nm) is approximately one-fifth of that of PhC_IC (2,700 nm) as determined by scanning electron microscopy (SEM) (Figure 1d and Supplementary Figure 2).

### Time course and Temperature dependency of the PhC_CF formation

To clarify the time dependency of the **PhC_CF** formation, the crystallization reaction induced by CFPC was monitored at 0.5 h, 1 h, 1.5 h, 2 h, 4 h, 6 h, 12 h, and 24 h with SEM. When PhM was expressed at 20 °C, the cubic crystals were first observed 2 h after the initiation of the translation reaction (Figure 2). The average sizes of the crystals measured after 2 h, 4 h, 6 h, 12 h, and 24 h were found to be 340 nm, 400 nm, 400 nm, 470 nm, and 580 nm, respectively. There were no crystals observed by SEM after 1.5 h of the reaction. When the expression of PhM was confirmed by SDS-PAGE at 0.5 h, 1 h, 1.5 h, 2.0 h, 3.0 h, and 4.0 h after the translation reaction at 20 °C, a band corresponding to PhM was observed after 1h, but the band was not observed at 0.5 h (Supplementary Figure 3a). These results indicate that insufficient amounts of PhM for the crystallization are obtained after 1.5 h.

**Figure 2.**
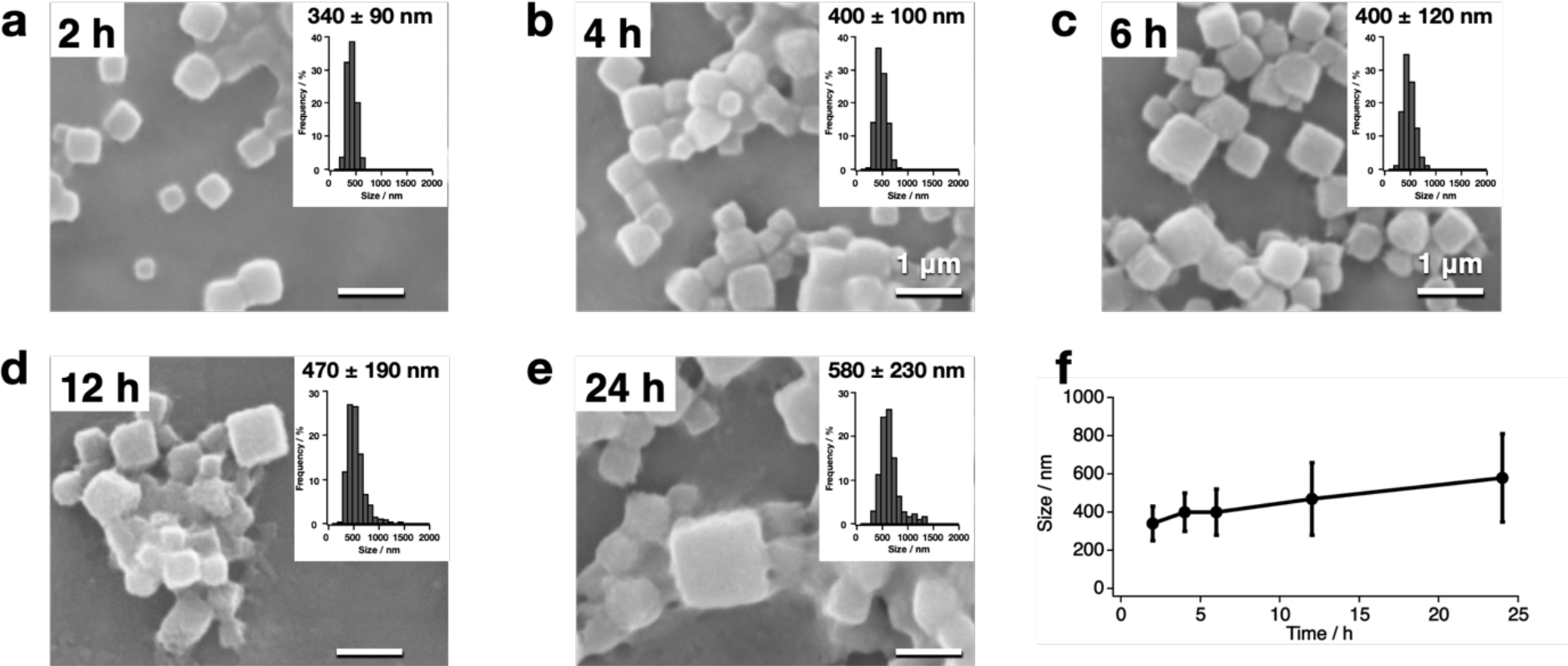
Time-dependent CFPC of **PhC_CF**. SEM images and size histograms of the purified **PhC_CF** after translation at 20 °C for (a) 2 h, (b) 4 h, (c) 6 h, (d) 12 h, and (e) 24 h. (f) Crystal size of purified **PhC_CF** over time. Scale bars = 1 μm .

To evaluate the temperature dependency of the **PhC_CF** formation, the translation reactions were performed at various temperatures. After 24 h, the crystals were formed with average sizes of 330 nm, 390 nm, 450 nm, 580 nm, and 1,170 nm at 4 °C, 10 °C, 15 °C, 20 °C, and 25 °C, respectively (Figure 3). At 15 °C and 25 °C, cubic crystals were observed, but many round crystals were observed at 4 °C and 10 °C. Although temperature is expected to affect the expression yield during translation reaction, the yields of PhM after 24 h did not differ significantly in the temperature range as confirmed by SDS-PAGE (Supplementary Figure 3b)^19^. Therefore, the crystal size and morphology are expected to be affected by the crystallization rate at various temperatures.

**Figure 3.**
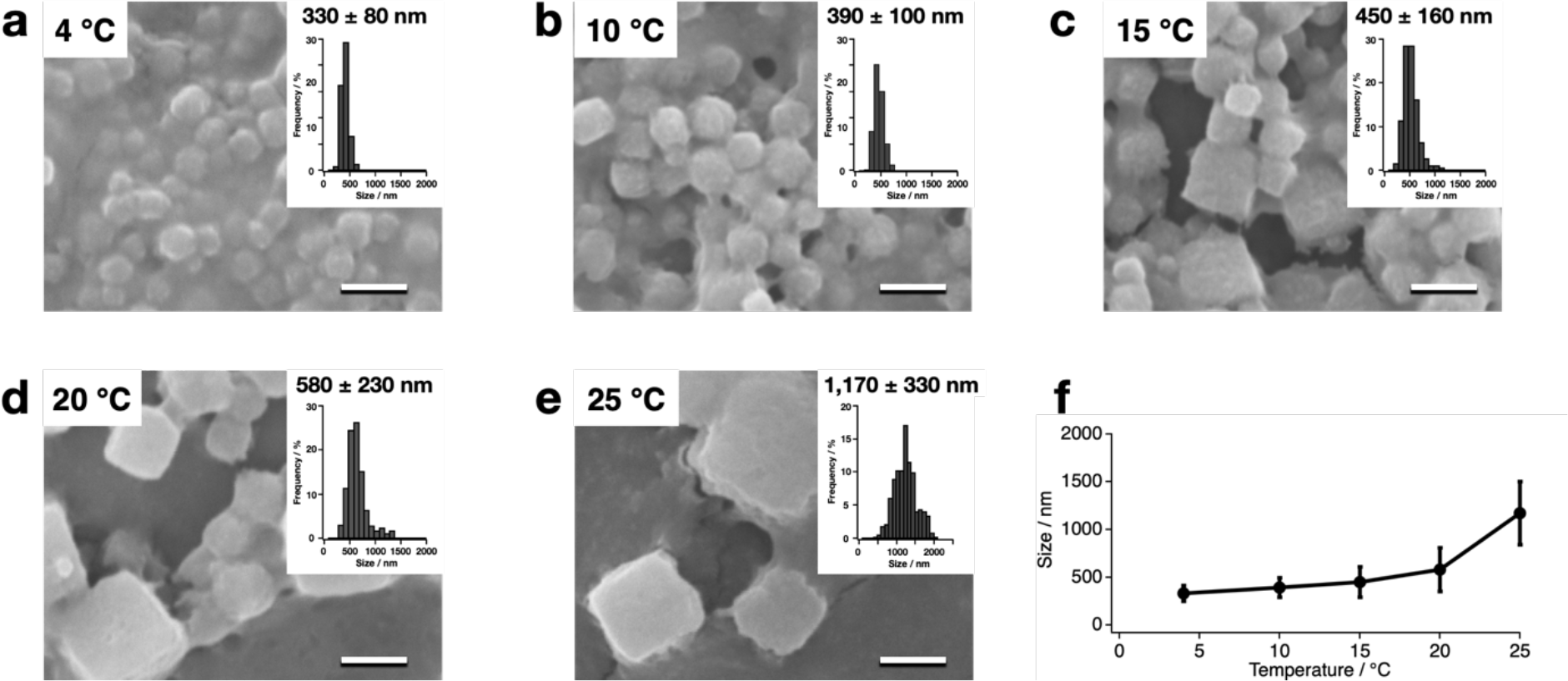
Temperature-dependent CFPC of **PhC_CF**. SEM images and size histograms of the purified **PhC_CF** after translation at (a) 4 °C, (b) 10°C, (c) 15°C, (d) 20 °C, and (e) 25 °C for 24 h. (f) Crystal size of purified **PhC_CF**s over temperature. Scale bars = 1 μm.

### Structural analysis of the nano-sized PhC_CFs

To collect diffraction data from the nano-sized **PhC_CFs**, the crystals isolated from the reaction mixture were diffracted using the micro-X-ray beam of the BL32XU beamline equipped with Serial Synchrotron Rotation Crystallography (SS-ROX) at SPring-8^26, 27^. **PhC_CF** obtained after 24h at 20°C (**PhC_CF_20°C/24h_**) was refined with a resolution of 1.80 Å, and has a space group (*I*23) and lattice parameters which are essentially identical to those of PhC_IC (PDB ID: 5gqm) (Supplementary Table 1). The root mean square deviation (RMSD) value of the Cα atoms for the structure from PhC_IC is 0.09 Å (Figure 4a). The main difference between **PhC_CF_20°C/24h_** and PhC_IC is that **PhC_CF_20°C/24h_** shows no electron density of nucleotide triphosphates (NTPs) bound to the monomer interface, which are observed in PhC_IC (Figure 4b-4e). The average *B*-factor values per residues of all atoms in **PhC_CF_20°C/24h_** show a large value for His76 because of the lack of cytosine triphosphate (CTP) interacting with His76 in PhC_IC (Figure 4b, 4d, and Supplementary Figure 4). In PhC_IC, the amino acid residues surrounding guanine triphosphate (GTP) and adenosine triphosphate (ATP) interact with the NTPs, but in **PhC_CF_20°C/24h_**, which lacks these electron densities, there is no significant difference in the side chain conformation between **PhC_CF_20°C/24h_** and PhC_IC (Figure 4b–4e). These results indicate that the NTP binding is not essential for the crystallization of PhM.

**Table 1.** Crystal size and Crystallographic data of **PhC_CF**.

| Crystals | Temperature (°C) | Reaction Time (h) | Crystal size (nm) | Resolution (Å) |
| --- | --- | --- | --- | --- |
| **PhC_CF<sub>20°C/1h</sub>** | 20 | 1 | —<sup>a</sup> | —<sup>b</sup> |
| **PhC_CF<sub>20°C/2h</sub>** | 20 | 2 | 340±90 | —<sup>b</sup> |
| **PhC_CF<sub>20°C/4h</sub>** | 20 | 4 | 400±100 | —<sup>b</sup> |
| **PhC_CF<sub>20°C/6h</sub>** | 20 | 6 | 400±120 | 2.50 |
| **PhC_CF<sub>20°C/12h</sub>** | 20 | 12 | 470±190 | 2.18 |
| **PhC_CF<sub>20°C/24h</sub>** | 20 | 24 | 580±230 | 1.80 |
| **PhC_CF<sub>4°C/24h</sub>** | 4 | 24 | 330±80 | —<sup>b</sup> |
| **PhC_CF<sub>10°C/24h</sub>** | 10 | 24 | 390±100 | —<sup>b</sup> |
| **PhC_CF<sub>15°C/24h</sub>** | 15 | 24 | 450±160 | 2.20 |
| **PhC_CF<sub>25°C/24h</sub>** | 25 | 24 | 1,170±330 | 1.87 |
| **PhC_IC** | 27 | 72 | 2,710±870<sup>c</sup> | 1.70<sup>d</sup> |

<sup>a</sup> no crystals observed by SEM. <sup>b</sup> no data sets were obtained. <sup>c</sup> Supplementary Figure 2. <sup>d</sup> reference 28.

**Figure 4.**
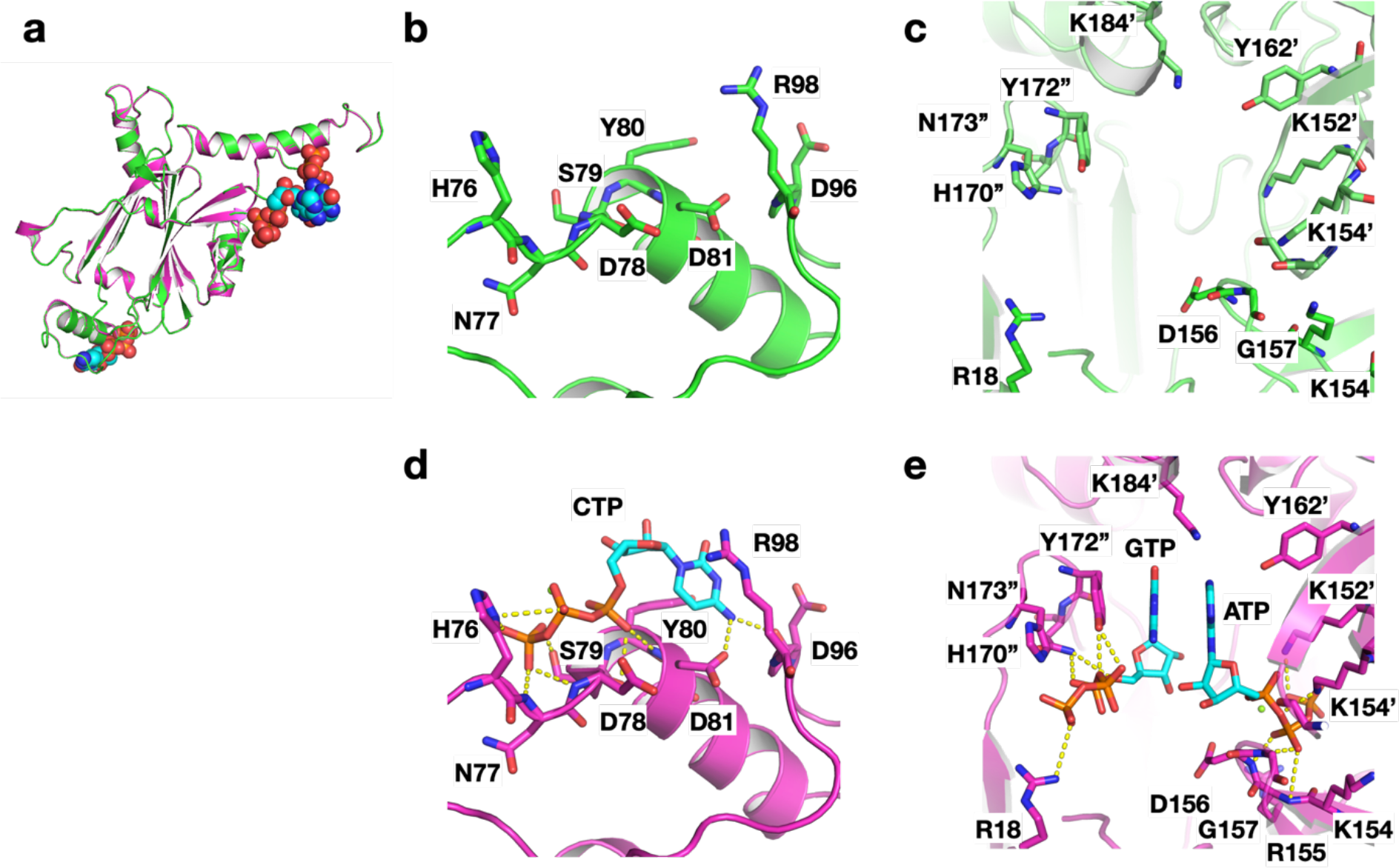
Comparison of crystal structure between **PhC_CF_20°C/24h_** and PhC_IC (PDB ID: 5gqm)^28^. (a) Superimposed structures of **PhC_CF_20°C/24h_** (green) and PhC_IC (magenta). (b and c) Close-up views of (b) CTP and (c) ATP/GTP binding sites in **PhC_CF_20°C/24h_**, (d and e) Close-up views of (d) CTP and (e) ATP/GTP binding sites in PhC_IC.

The crystals formed with CFPC under various conditions provided data sets suitable for structural analysis (Table 1, Supplementary Table 1, and 2). The crystal structures were refined with a resolution range of 1.80-2.50 Å. The RMSD values of the Cα atoms from PhC_IC are less than 0.29 Å, indicating that the overall structure of PhC_IC is retained in **PhC_CFs**. While no data sets were obtained for the crystals formed at 20 °C for 2 h and 4 h, and at 4 °C and 10 °C for 24 h due to fewer indexed images, we obtained a data set for **PhC_CF_20°C/6h_** with a resolution of 2.50 Å (Table1, Supplementary Table 1, and 2). These results show that the CFPC reaction of PhC under optimized conditions successfully produces nanocrystals with sufficient quality to obtain a high-resolution structure in only 6 hours. This reaction time is dramatically reduced from the cultivation time (>3 days) required to obtain comparable high-quality crystals using insect cells^15^.

### Determination of the structure of CipA by CFPC

After successfully validating CFPC in crystallizing PhC, we applied this method to overcome current challenges in structure determination of CipA since it provides an opportunity to add chemical compounds during crystallization. Crystalline inclusion protein A (CipA), a hydrophobic protein of 104 amino acid residues, spontaneously forms crystalline aggregates in *Photorhabdus luminescens*, an entomopathogenic bacterium^29^. Its native function is postulated to be involved in nematode symbiosis or pathogenesis^30^. It can also form a crystallized aggregation in *E.coli*, which is used as a template for constructing solid-nanomaterials. Its structure has not yet been determined^22^. CipA crystals formed in *E. coli* (CipAC_EC) with an average size of 410 ± 80 nm were diffracted with a resolution of 2.8 Å using the SS-ROX method at the BL32XU beamline of SPring-8 (Supplementary Figure 5a, 5b, Supplementary Table 3, and 4). However, we could not determine the structure with a high R-value when the predicted structure using AlphaFold2 (AF2) was used as the initial model^31^. This is because of the large twin fraction of 0.42 (Supplementary Table 3). Next, we attempted to determine the structure of CipA by reducing the twin fraction using CFPC. When CipA was expressed by CFPC with dialysis method^32^, a white precipitate appeared in the solution mixture after 24 h (Supplementary Figure 5c). The SEM image and the MALDI TOF-MS of the precipitate showed that the precipitate is the CipA crystal (**CipAC_CF**) with an average size of 3,400 ± 880 nm (Supplementary Figure 5d and 5e). This is eight times larger than the size of CipAC_EC. Structural analysis of **CipAC_CF** was attempted with data of a resolution of 1.61 Å obtained using the small wedge method at BL32XU of SPring-8^27^. However, the structure could not be determined because the twin fraction was still too high (0.42), as it is in *E.coli*. To overcome the twinning issue occurring in the high quality crystals, we recognized that the CFPC method permits addition of reagents which inhibit twinning, such as ethanol, 1,4-dioxane, PEGs, Dextran, and TEG^33, 34^. The SEM images show that the **CipAC_CFs** crystallized in the presence of additives have slightly larger or similar sizes and similar shapes relative to **CipAC_CF** crystallized in the absence of additives (Supplementary Figure 6). This indicates that the CFPC method can be expanded to include various chemical manipulations to crystals, as well as addition of chemical compounds and proteins to crystals and proteins during the crystal growth process.

X-ray diffraction experiments of the crystals showed that **CipAC_CF** crystallized in the presence of 3 v/v % 1,4-dioxane dramatically reduces the twin fraction to 0.10, with a resolution of 2.11 Å (Supplementary Table 3 and 4). The crystal is tetragonal, with a *I*4 space group having unit cell parameters *a*=*b*=60.1 Å, *c* = 54.0 Å, α = β = γ = 90° (Supplementary Table 3 and 4). The structure was determined by the molecular replacement method using the search model created by AF2^31^ *via* ColabFold^35^. The structure of the CipA monomer consists of the N-terminal arm followed by three β-strands β1, β2, β3, α-helix, and β-strands β4 and β5. Except for the N-terminal arm, the globular domain is a typical oligonucleotide /oligosaccharide-binding (OB) fold (Figure 5a). In the crystal lattice, the four α-helices from each monomer form a four-helix bundle around the crystallographic 4-fold symmetry axis and exist as a tetramer. This tetramer is consistent with the results of PISA^36^ prediction of oligomeric states and is considered the basic unit of crystal growth. As for the interactions between monomers in the tetramer, hydrogen bonds (N_δ_/Asn62_i_–O/Leu59_ii_, N_δ_/Asn62_i_–O/Leu60_ii_, N_δ_/Asn62_i_– O_δ_/Asn62_ii_, O/Ala66_i_–O_ζ_/Tyr68_ii_, and O/Met104_i_–N_ζ_/Lys78_ii_) are formed between each helix (Figure 5b). In addition, the β1 and β5 strands of the neighboring molecule form a new β-sheet with hydrogen bonds (N_ζ_/Lys56_i_–O_δ_/Asp34_ii_, N_δ_/Asn98_i_–O_δ_/Asp34_ii_, N/Val100_i_–O/Val31_ii_, O/Val100_i_–N/Val31_ii_, N/Ile102_i_–O/Met29_ii_, N/Ile102_i_–N/Met29_ii_, and N/Met104_i_–O/Ile27_ii_) and salt bridge (N_ζ_/Lys56_i_–O_δ_/Asp34_ii_) (Figure 5b). The interaction between the edges of each tetramer forms the basic crystal lattice, and it is further stabilized by the embedded N-terminal arm of the neighboring monomer molecule at the cleft created between the tetramer-tetramer interface (N/Asp4_i_-O_δ_Asp61_ii_, O/Asp4_i_-N/Asn58_ii_, O_δ_/Asp4_i_-N_δ_/Asn58_ii_, N/His6_i_-O/Lys56_ii_, N_ε_/His6_i_-O/Met37_ii_, O_γ_/Ser8_i_-O_ζ_/Tyr96_ii_, and O/Ser8_i_-O_ζ_/Tyr96_ii_) (Figure 5c).

**Figure 5.**
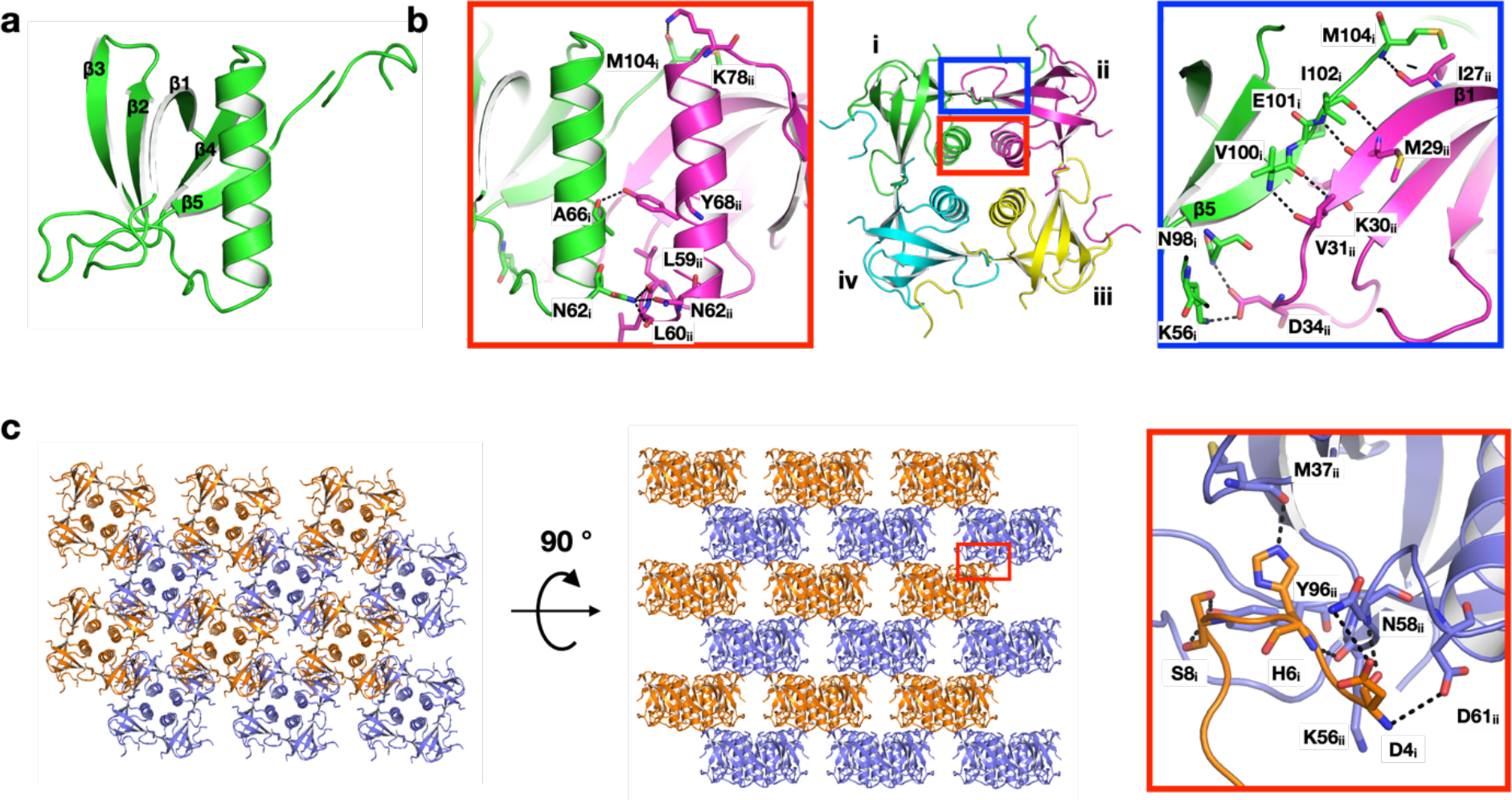
Crystal structure of **CipAC-CF** with 1,4-dioxane. The structure of (a) monomer and (b) tetramer. (a) CipA monomer are consists of the N-terminal arm followed by three β-strands β1, β2, β3, α-helix, and β-strands β4 and β5. (b) the interactions between monomers (i, ii, iii and iv) in the tetramer. (c) Lattice structure and interactions between tetramers. Hydrogen bonds are indicated with black dotted lines.

A search of the Dali server^37^ search for similar structures shows, as expected, OB-fold domain-containing proteins. Among them, a pentameric B subunit of heat-labile enterotoxin type IIB (PDB ID: 1QB5) from a pathogenic bacterium with high structural similarity (z-score >9.0) was found. Even though the sequence homology was less than 10%, the topology of the monomers of this protein and CipA is very similar, with an RMSD value of 2.2 Å in equivalent Cα atoms. Although these proteins have different oligomerization states, each monomer forms a bundle of α-helices around the central symmetry axis, and the β-strands at adjacent monomers form β-sheets at the outer rim of the complex (Supplementary Figure 7).

## Discussion

We have succeeded in determining crystal structures of two types of proteins with high-resolution using CFPC. ICPC of both PhC and CipA are believed to be influenced by the complex environment of living cells^5, 22^. However, a comparison of the structures of PhCs by ICPC and CFPC suggests that the encapsulation of NTPs in PhCs is not essential for the crystallization. For the crystallization of CipA, the formation of the twin crystal by ICPC could be inhibited by the addition of foreign molecules in CFPC to obtain crystals used for the structural analysis. These results suggest that strict cellular environments are not required for their crystallization. Thus, by using CFPC, we were able to determine the crystallization factors in cell environments, which were difficult to investigate by ICPC alone. In addition, the structures of some in-cell protein crystals have been determined by *in vitro* crystallization after purification^4, 38^. In other words, proteins that can be crystallized *in vitro* are candidates for using CFPC to obtain crystals. These results indicate that there are opportunities to use CFPC to obtain crystals of other recombinant proteins.

Protein crystals with high-resolution diffraction data were obtained from small reaction volumes and short CFPC timeframes. CFPC of PhC allowed us to obtain the 2.50 Å structure for crystals obtained only 6 hours after initiating the reaction. ICPC using insect cells requires three days to obtain PhC_IC after virus infection (Table 1)^15^. This is because insect cells involve various cellular processes in producing the target protein. The CFPS reaction system is dedicated to producing the target protein and the crystals. Furthermore, CFPC using cell extracts allows crystal formation independent of the reaction scale. The reaction was carried out by dialyzing a mixture of WEPRO^®^7240 (5 μL), the mRNA solution (5 μL), and 10 μL SUB-AMIX^®^ SGC against 1.0 mL SUB-AMIX^®^ SGC. After 24 hours at 20 °C, crystals were collected. The average size of the crystals was 610±150 nm, which is essentially identical to the average size of **PhC_CF_20°C/24h_** (Supplementary Figure 8). Thus, it was found that CFPC can efficiently synthesize high-quality protein crystals by taking advantage of the smaller reaction scale. This provides a solution to the problem of low yields of high-quality crystals produced by ICPC and shows great potential to facilitate crystal screening, which is not feasible using previously reported methods and ICPC.

The crystal structure of CipA was determined at high resolution by adding chemical reagents to the CFPC reaction mixture. **CipAC_CF** forms a multilayer structure composed of the tetramers as the building blocks (Figure 5). In addition to 1,4-dioxane, which is known to be a twin inhibitor^39^, PEGs were found to reduce the twin fraction (Supplementary Table 3). In particular, a significant reduction by PEG400 was observed. The effect is presumed to be similar to the effect provided by 2-methyl-2,4-pentanediol (MPD), rather than the exclusion volume effect of PEG with a large molecular weight^33^. Therefore, twin formation may be inhibited by binding of PEG to hydrophobic patches on the protein surface and filling the voids. On the other hand, CipAC_EC prepared by ICPC gave high-resolution diffraction data but the structure could not be determined due to the high twin fraction (Supplementary Table 3). In *E. coli*, a lack of interacting molecules, such as PEG400, on the surface of CipA may lead to incorporation of inverted tetramers into the multilayered crystal structure of CipA_EC, resulting in a large twin fraction (Supplementary Figure 6). In this way, CFPC permits control of the crystallization process to obtain high quality protein crystals by adding chemical reagents. CFPC is useful for investigating crystallization steps in living cells. Several model proteins have been used to study the ICPC mechanism^12^. These model proteins can be crystallized even in an impurity-rich intracellular environment. These studies suggest that abundant intracellular and organelle endogenous proteins play roles as precipitants or crowding agents, like polyethylene glycols. Furthermore, ICPC has been attempted by adding a crystallization reagent, but no characteristic effect on the crystallization has been confirmed^12^. Since the intracellular reactions induced by ICPC are complex, it remains challenging to identify common crystallization factors for model proteins. In this study, the addition of PEG was found to inhibit twinning and promote the formation of high-quality crystals with a larger size (Supplementary Figure 6). It has been reported that PEG induces liquid-liquid phase separation (LLPS) in the reaction of CFPS^40^. Furthermore, LLPS plays an important role during *in vitro* protein crystallization^41^. Thus, it is inferred that the LLPS is one factor involved in promoting the crystallization of polyhedra and CipA in the CFPC solution. A detailed study of this proposal is underway.

## Conclusion

We established a CFPC method to rapidly obtain protein crystals in microliter volumes within a few hours without complicated purification and crystallization procedures. Furthermore, high-resolution structures of proteins were obtained using the nanocrystals. Although ICPC has been expected to become an important tool in crystal structure analysis, crucial challenges remain because the crystals are not formed in suitable amounts and quality to provide high-resolution structures. We used CFPC to enable rapid screening of reaction conditions such as temperature, time, and the effects of additives and achieved preparation of high-quality protein crystals suitable for structure analysis. These results indicate that CFPC, a hybrid method of ICPC and *in vitro* protein crystallization, will likely be a powerful HTS tool for crystal structure analysis of unstable, low-yield, and substrate-binding proteins which are currently considered challenging to analyze using conventional protein crystallization methods.

## Acknowledgements

We thank the Suzukakedai Analysis Division and Technical Department, Biomaterials Analysis Division, Tokyo Institute of Technology for supporting us with the corresponding measurements. This work was supported by JSPS KAKENHI Grant Nos. JP19H02830, Grant-in-Aid for Scientific Research on Innovative Areas “Molecular Engines” (JP18H05421) to T.U. and JP18K05140 to S.A., and Adaptable and Seamless Technology Transfer Program through Target-driven R&D (JPMJTR20U1) from the Japan Science and Technology Agency to T.U. Synchrotron radiation experiments were conducted under the approval of 2019B2561, 2020A2541, 2021A2541, and 2021B2544 at SPring-8.

## Author contributions

S.A. and T.U. designed the research. S.A., J. T., M. K., S. K., K. H., K. Y. and A. K. carried out the experiments. All authors analyzed the data, discussed the results and co-wrote the paper.

## Accession Codes

Atomic coordinate of **PhC_CF_20°C/24h_** and **CipAC** crystallized with 3 % 1, 4-dioxane have been deposited in the Protein Data Bank under the code of 7XHR and 7XHS, respectively.

