## Supplemental Information for "Cell-free Protein Crystallization for Nanocrystal Structure Determination"

### Table of Contents

**Methods**

Materials 3

Scanning Electron Microscopy (SEM) 3

Cell-free synthesis and crystallization of PhM 3

X-ray crystal structure analysis of PhC_CFs 3

Cell-free synthesis and crystallization of CipA 4

X-ray crystal structure analysis of the CipACs 5

**Supplementary Figures 1-8** 6-13

**Supplementary Tables 1-4** 14-17

**Supplementary References** 18

### Methods

#### Materials

All reagents were purchased from TCI, Wako, Nacalai Tesque, Sigma–Aldrich, and Life Technologies and were used without further purification.

#### Scanning Electron Microscopy (SEM)

The morphologies of purified crystals were confirmed by scanning electron microscopy (SEM). After substituting PBS with Milli-Q water, the crystals were dried and observed by SEM. SEM analyses were performed on JCM-6000 Neoscope (JEOL).

#### Cell-free synthesis and crystallization of PhM

Expression and crystallization of PhM were performed using a WEPRO7240 Expression Kit (CellFree Sciences). The gene of polyhedrin was cloned into the PEU-E01-MCS vector (CellFree Sciences) for the polyhedrin expression. The plasmid was amplified in DH5α bacteria and purified using the Qiagen Plasmid Midi Kit. Transcription was performed in 1.5 mL tubes according to the protocol of the expression kit. After incubation for 6 h at 37 °C, mRNA was used for translation. Translation reactions were carried out using the bilayer method. 20 μL of reaction mixture containing 10 μL of WEPRO7240 wheat germ extract, 10 μL of prepared mRNA, 40 mg/mL creatine kinase was overlaid with 200 μL of SUB-AMIX SGC solution in a microtube and incubated at 20 °C for 24 h. The crystals were collected by centrifuge and washed with PBS several times. Time-dependent and temperature-dependent crystallization were performed by the same method except for the change of time and temperature.

#### X-ray crystal structure analysis of PhC_CFs

Before the data collection, the crystals were immersed in PBS buffer containing 50 % ethylene glycol and then loaded onto the MicroLoops (Mitegen), followed by frozen in liquid nitrogen. The data diffraction of PhCs was collected at 100K using the beamline of BL32XU at SPring-8 with an X-ray wavelength of

1.00Å. The whole data collection was automated by *ZOO* system, including sample change by a robot[1]. Serial Synchrotron Rotation Crystallography (SS-ROX) method, which was developed to collect diffraction data of microcrystals efficiently, was employed[2, 3]. A microbeam of 1.2 μm (vertical) x 1.0 μm (horizontal) was used. The datasets were collected using a helical rotation per image of 0.25°/1μmand a frame rate of 58.824 Hz (~2 x $10^{10}$ photons/frame). The index was performed using *CrystFEL* version 0.6.3[4] with *Dirax*[5] and *Mosflm*[6]. The number of indexed images for **PhC_CF$_{20°C/6h}$**, **PhC_CF$_{20°C/12h}$**, **PhC_CF$_{20°C/24h}$**, **PhC_CF$_{15°C/24h}$**, and **PhC_CF$_{25°C/24h}$** are 1531, 8754, 12846, 3834, and 12376, respectively. Integrated intensities were merged by process_hkl in the *CrystFEL* suite. The datasets were not obtained for **PhC_CF$_{20°C/2h}$**, **PhC_CF$_{20°C/4h}$**, **PhC_CF$_{4°C/24h}$**, and **PhC_CF$_{10°C/4h}$** due to fewer indexed images (1, 235, 0, and 258, respectively) (Supplementary Table 1). The structure of **PhC_CF$_{20°C/24h}$** was solved by rigid-body refinement with *phenix.refine*[7] using the previously determined structure (PDB ID: 5GQM). The refinement process of the protein structure was performed using *phenix.refine* and *REFMAC5* in the *CCP4* suite[7, 8]. Rebuilding was completed using *COOT*[9] based on sigma-A weighted (2*Fo*-*Fc*) and (*Fo*-*Fc*) electron density maps. The models were subjected to quality analysis during the various refinement stages with omit maps and *RAMPAGE*[10].

**Cell-free synthesis and crystallization of CipA**

Expression and crystallization of CipA were performed with dialysis method using a WEPRO7240 Expression Kit (CellFree Sciences). Transcription reaction was performed with the same as PhC_CF. Translation reactions were carried out using the dialysis method. The reaction mixture containing 20 μL of WEPRO7240 wheat germ extract including 40 mg/mL creatine kinase, 20 μL of prepared mRNA, and 40 μL of SUB-AMIX SGC solution was dialyzed against 2.5 mL SUB-AMIX SGC solution at 20 °C for 72 h. As for crystallization of CipA with additives, the reaction mixture containing 16 μL of WEPRO7240 wheat germ extract including 40 mg/mL creatine kinase, 16 μL of prepared mRNA, 16 μL additive of appropriate concentration, and 32 μL of SUB-AMIX SGC solution was dialyzed against 2.5 mL SUB-

AMIX SGC solution including additives at 20 °C for 72 h. The crystals were collected by centrifuge and washed PBS several times.

**X-ray crystal structure analysis of the CipACs**

Before the data collection, the crystals were immersed in PBS buffer containing 50 % ethylene glycol and then loaded onto the MicroLoops (Mitegen), followed by frozen in liquid nitrogen. The diffraction data of CipACs were collected at 100K using the beamline of BL32XU at SPring-8 with an X-ray wavelength of 1.00Å. The crystal positions in a cryo-loop were identified by a low-dose raster scan. The complete datasets were obtained by merging multiple small-wedge (10° each) datasets collected from single crystals. Collected datasets were automatically processed and merged by *KAMO*[11]. Each dataset was indexed and integrated using *XDS*[12]. The datasets were subjected to hierarchical clustering by a pairwise correlation coefficient of intensities. The datasets in each cluster were scaled and merged using outlier rejections implemented in *KAMO*. The groups with the highest $CC_{1/2}$ were chosen for downstream analyses. Twinning was analyzed with *phenix.xtriage*[13]. The structure was solved by molecular replacement with *Phaser-MR*[13] of *PHENIX* suite using the predicted structure by AlphaFold2[14]. Rebuilding was completed using *COOT*[9] based on sigma-A weighted (2*Fo*-*Fc*) and (*Fo*-*Fc*) electron density maps. The models were subjected to quality analysis during the various refinement stages with omit maps and *RAMPAGE*[10].

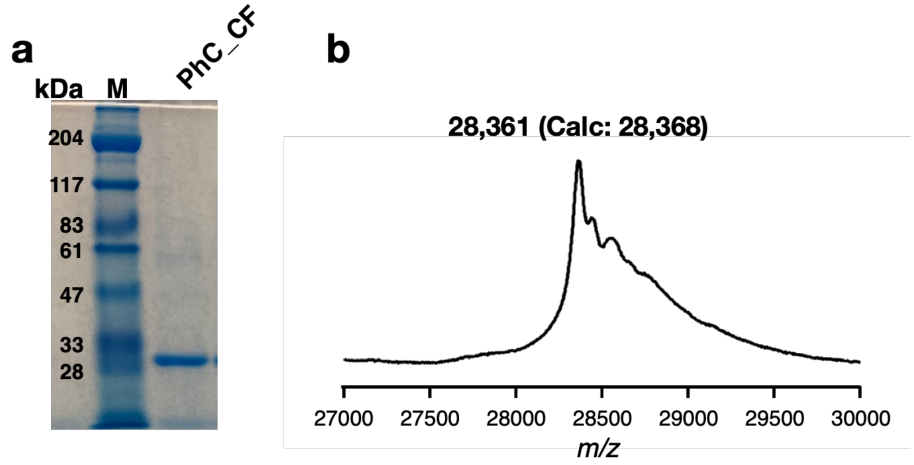


**Supplementary Figure 1.** (a) SDS-PAGE analysis and (b) MALDI TOF-MS of purified **PhC_CF**.

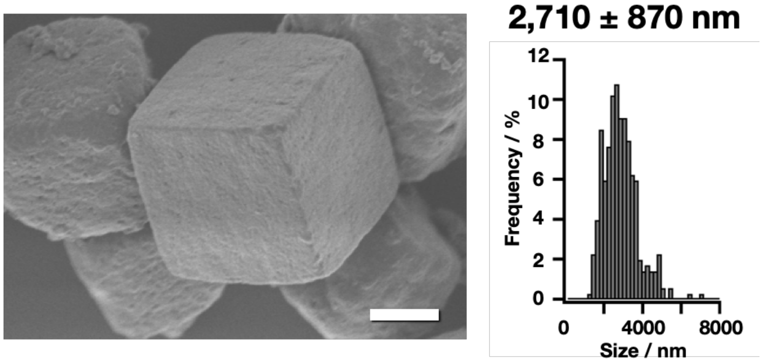


**Supplementary Figure 2.** SEM image and size histogram of purified PhC_IC. Scale bar = 1 μm.

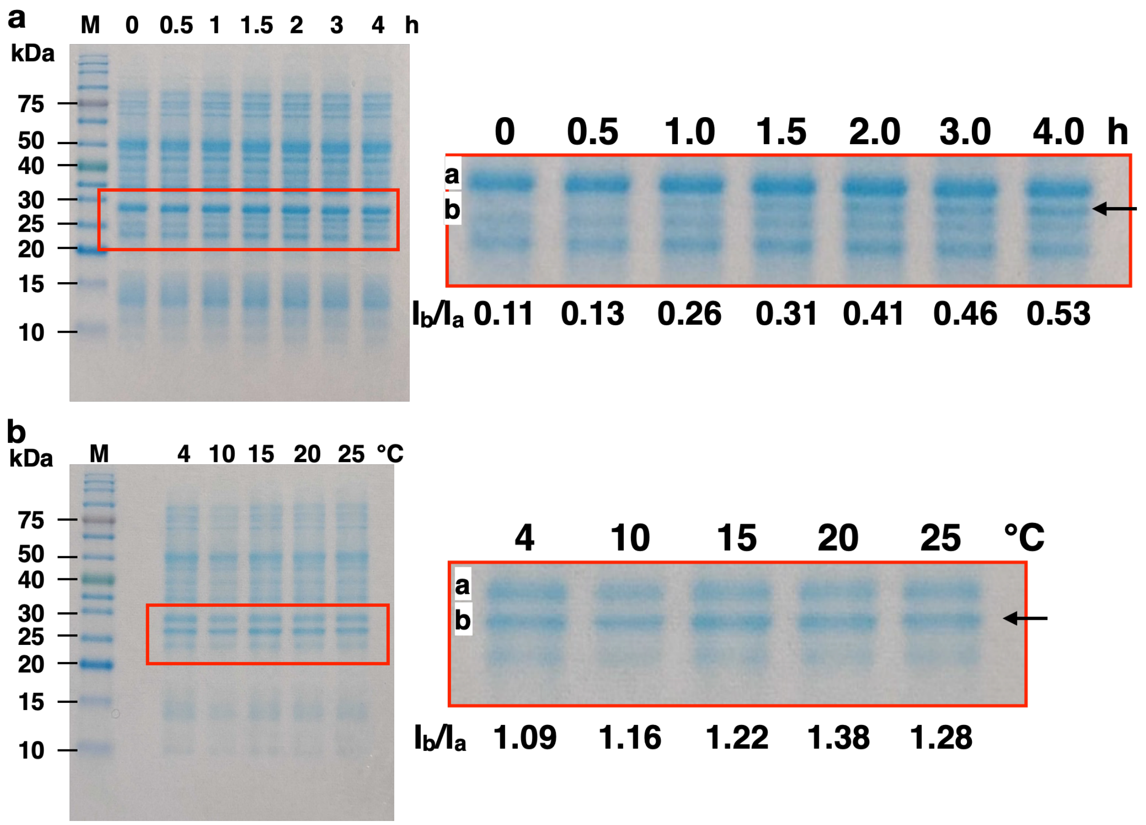


**Supplementary Figure 3.** SDS-PAGE analysis of CFPC of PhM. (a) Time dependence and (b) temperature dependence of PhM expression. The intensities of the bands are measured with Image J. The ratio of the intensity (Ib/Ia) of band b corresponding to PhM to that of the band a is shown.

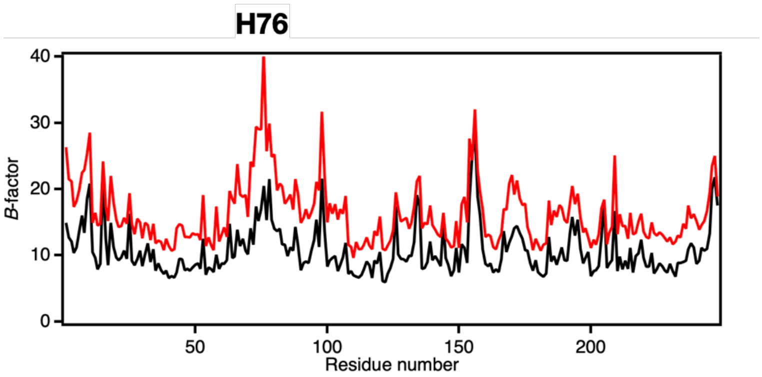


**Supplementary Figure 4.** Average *B*-factor values per residues of all atoms in **PhC_CF$_{\mathbf{20°C/24h}}$** (red line) and PhC_IC (black line).

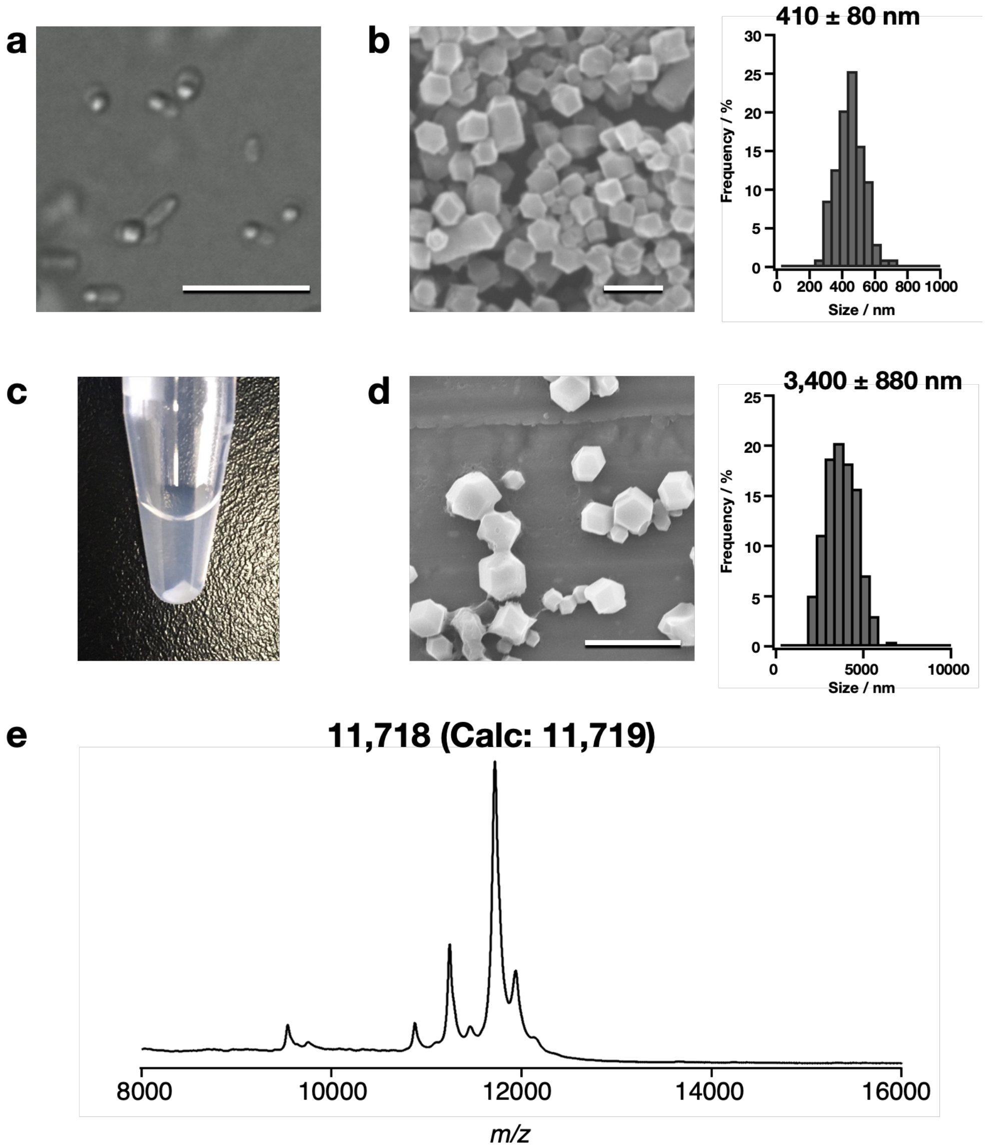


**Supplementary Figure 5.** (a) *E.coli* produced CipAC. Scale bar = 10 μm. (b) SEM image and size histogram of purified CipAC_EC. Scale bar = 1 μm. (c) Photograph of the tube after CFPC of CipA. (d) SEM image and size histograms of purified **CipAC_CF**. Scale bar = 10 μm. (e) MALDI TOF-MS of purified **CipAC_CF**.

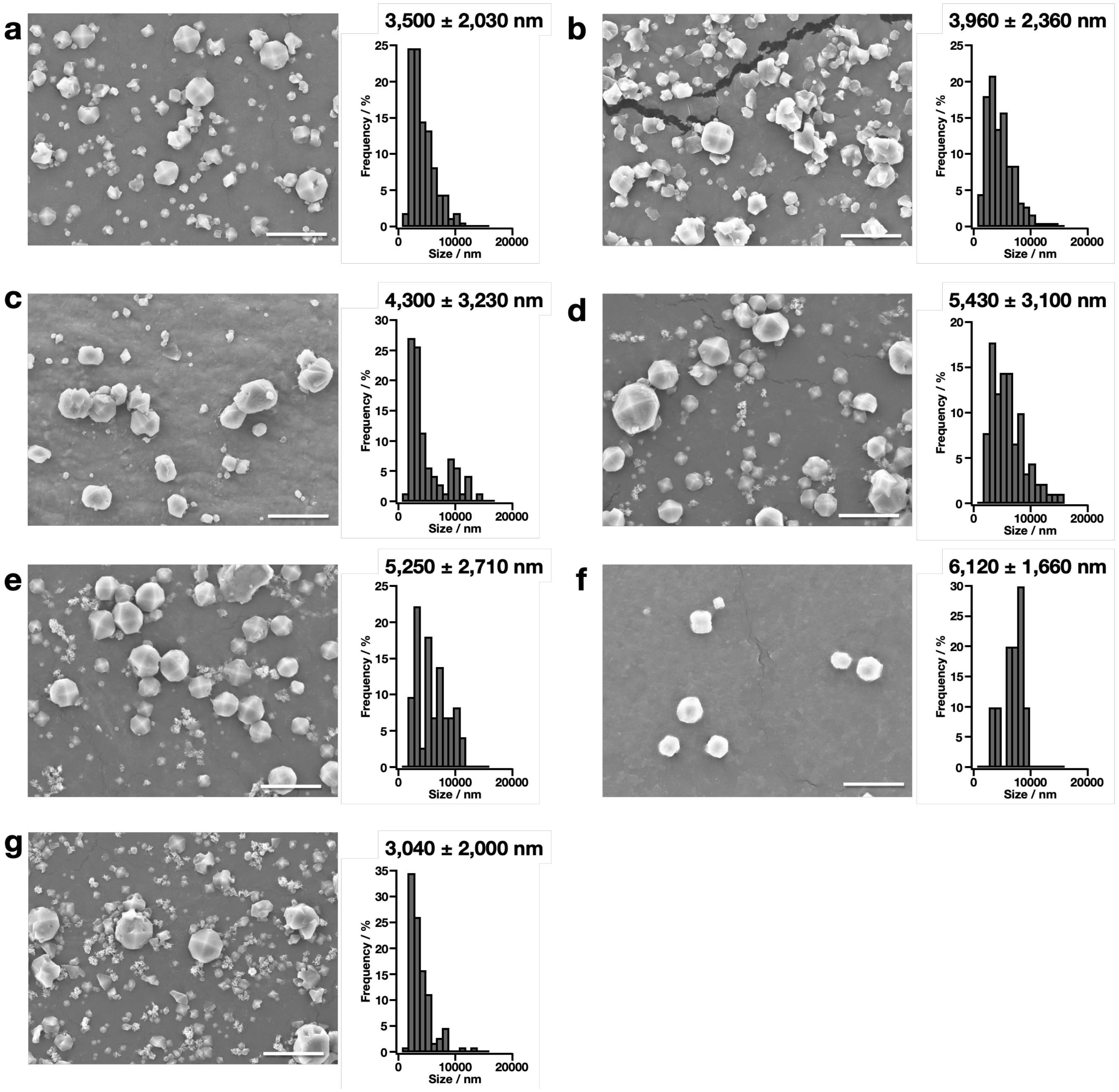


**Supplementary Figure 6.** SEM images of purified **CipAC_CFs** with additives. (a) 3 v/v % EtOH, (b) 3 v/v% Dioxane, (c) 1 v/v% PEG400, (d) 1 w/v % PEG3350, (e) 1 w/v % PEG8000, (f) 2 w/v % Dextran, (g) 1 v/v% TEG. Scale bars = 20 μm

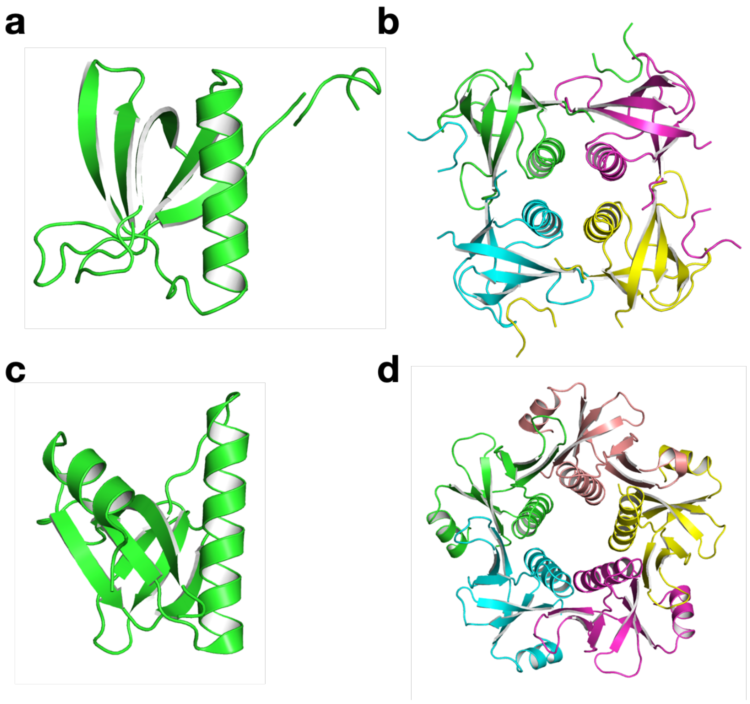


**Supplementary Figure 7.** Structure comparison of (a and b) CipA and (c and d) heat-labile enterotoxin type IIB (PDB ID: 1QB5).

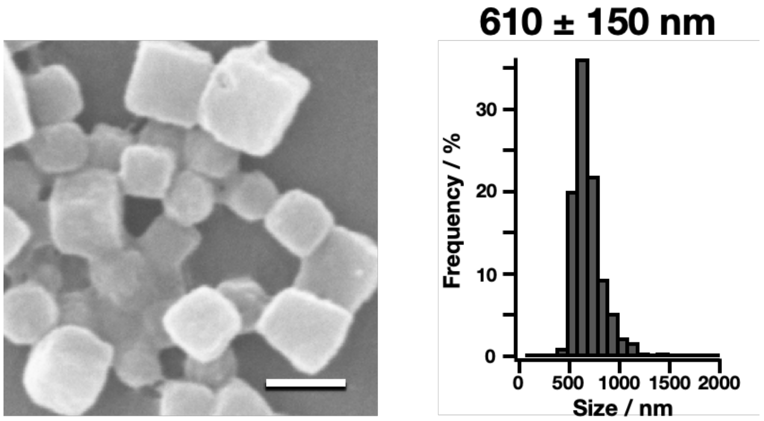


**Supplementary Figure 8.** SEM image and size histogram of **PhC_CF** produced by dialysis with 20 μL reaction scale. Scale bar = 1 μm

**Supplementary Table 1.** Crystallographic data of **PhC_CF**.

| | 20 °C/2h | 20 °C/4h | 20 °C/6h | 20 °C/12h | 20 °C/24h | 4 °C/24h | 10 °C/24h | 15 °C/24h | 25 °C/24h |
|---|---|---|---|---|---|---|---|---|---|
| No. of loops | 4 | 4 | 4 | 4 | 4 | 4 | 4 | 4 | 4 |
| No. of indexed images | 1 | 91 | 1531 | 8754 | 12846 | 0 | 258 | 3834 | 3807 |
| Space group | | | *I*23 | *I*23 | *I*23 | | | *I*23 | *I*23 |
| Crystal cell (Å) $a = b = c$ | | | 104.3 | 103.3 | 104.4 | | | | |
| Resolution range (Å) | | | 50-2.50 (2.51-2.50) | 50-2.18 (2.19-2.18) | 50-1.80 (1.81-1.80) | | | 50-2.20 (2.21-2.20) | 50-1.87 (1.88-1.87) |
| Observations | | | 427,843 (6,586) | 343,3242 (59,651) | 6,907,468 (114,912) | | | 1,489,024 (27,270) | 2,048,341 (33158) |
| Unique reflections | | | 6,679 (148) | 9,832 (253) | 17,297 (414) | | | 9,786 (261) | 15,459 (367) |
| Completeness (%) | | | 100 (100) | 100 (100) | 100 (100) | | | 100 (100) | 100 (100) |
| Multiplicity | | | 64.1 (44.5) | 349 (236) | 399 (277) | | | 152.2 (105) | 133 (90) |
| I/σ | | | 3.8 (1.24) | 7.3 (1.28) | 5.5 (1.53) | | | 4.8 (1.14) | 4.5 (1.33) |
| $CC_{1/2}$ | | | 0.9457 (0.5054) | 0.98821 (0.5854) | 0.9899 (0.5326) | | | 0.9759 (0.4000) | 0.9696 (0.4921) |

Values in parentheses are for the highest-resolution shell.

**Supplementary Table 2.** Refinement statics of **PhC_CF**.

| | 20 °C/2h | 20 °C/4h | 20 °C/6h | 20 °C/12h | 20 °C/24h | 4 °C/24h | 10 °C/24h | 15 °C/24h | 25 °C/24h |
|---|---|---|---|---|---|---|---|---|---|
| Resolution range (Å) | - | - | 42.58-2.50 | 36.63-2.18 | 42.30-1.80 | - | - | 36.9-2.20 | 42.31-1.87 |
| Reflection used | | | 6,679 | 9,829 | 17,297 | | | 9,785 | 15,457 |
| R-factor (%) | | | 17.36 | 17.17 | 15.44 | | | 17.27 | 17.16 |
| Free R-factor (%) | | | 26.79 | 23.53 | 18.66 | | | 24.07 | 22.56 |
| R.m.s.deviations from ideal | | | | | | | | | |
| Bond length (Å) | | | 0.008 | 0.008 | 0.014 | | | 0.008 | 0.010 |
| Angle (°) | | | 1.12 | 0.96 | 1.77 | | | 0.95 | 1.06 |
| Ramachandran plot (%) | | | | | | | | | |
| Favored region | | | 96.73 | 97.96 | 97.96 | | | 96.73 | 97.55 |
| Allowed region | | | 3.27 | 2.04 | 2.04 | | | 3.27 | 2.45 |
| Outlier region | | | 0.0 | 0.0 | 0.0 | | | 0.0 | 0.0 |

**Supplementary Table 3.** Crystallographic data of CipAC_EC and CipAC_CF

| | CipAC_EC | **CipAC_CF** | | | | | | | |
|---|---|---|---|---|---|---|---|---|---|
| | | No additive | EtOH 3 v/v% | Dioxane 3 v/v% | PEG400 1 v/v % | PEG3350 1 w/v % | PEG8000 1 w/v % | Dextran 2 w/v% | TEG 1 v/v % |
| Space group | *I*4 | *I*4 | *I*4 | *I*4 | *I*4 | *I*4 | *I*4 | *I*4 | *I*4 |
| Crystal cell (Å) $a = b$, and $c$ | 61.2, 53.8 | 60.4, 53.2 | 61.1, 54.0 | 61.1, 54.0 | 61.3, 54.2 | 61.1, 54.0 | 61.1, 54.0 | 61.0, 54.0 | 61.2, 54.1 |
| Resolution range (Å) | 50-2.80 (2.81-2.80) | 50-1.61 (1.71-1.61) | 50-1.98 (2.10-1.98) | 50-2.11 (2.24-2.11) | 50-2.47 (2.62-2.47) | 50-2.38 (2.52-2.38) | 50-2.08 (2.21-2.08) | 50-1.98 (2.10-1.98) | 50-2.48 (2.63-2.48) |
| Observations | 94,065 (1,261) | 3,412,217 (480,752) | 338,736 (50,925) | 111,712 (18,248) | 46,092 (7,428) | 43,814 (7,064) | 253,733 (37,516) | 338,784 (52,079) | 68,170 (11,178) |
| Unique reflections | 2,504 (62) | 12,425 (2,025) | 6,991 (1,125) | 5,787 (935) | 3,662 (582) | 4,080 (641) | 6,050 (994) | 6,987 (1,140) | 3,595 (574) |
| Completeness (%) | 100 (100) | 100 (100) | 100 (100) | 100 (100) | 99.9 (100) | 99.9 (100) | 100 (100) | 100 (100) | 100 (100) |
| Multiplicity | 37.6 (20.3) | 274.5 (237.4) | 48.5 (45.3) | 19.3 (19.5) | 12.6 (12.8) | 10.7 (11.0) | 41.9 (37.7) | 48.5 (45.7) | 19.0 (19.5) |
| I/σ | 5.2 (1.2) | 25.39 (0.7) | 12.46 (1.6) | 9.04 (1.7) | 6.49 (1.5) | 7.09 (1.8) | 11.01 (1.6) | 13.22 (1.7) | 8.73 (1.6) |
| $CC_{1/2}$ | 0.9139 (0.317) | 0.999 (0.593) | 0.991 (0.543) | 0.994 (0.587) | 0.979 (0.499) | 0.979 (0.375) | 0.993 (0.526) | 0.992 (0.693) | 0.980 (0.589) |
| $<I^2>/<I>^2$ [a] | 1.565 | 1.537 | 1.584 | 1.893 | 1.954 | 1.751 | 1.741 | 1.592 | 1.732 |
| Twin fraction [a] | 0.42 | 0.42 | 0.38 | 0.10 | 0.14 | 0.28 | 0.22 | 0.36 | 0.30 |

[a] Twinning was analyzed with *phenix.xtriage*. Values in parentheses are for the highest-resolution shell.

**Supplementary Table 4.** Refinement statics of CipAC_EC and CipAC_CF

| | CipAC_EC | **CipAC_CF** | | | | | | | |
|---|---|---|---|---|---|---|---|---|---|
| | | No additive | EtOH 3 v/v% | Dioxane 3 v/v% | PEG400 1 v/v % | PEG3350 1 w/v % | PEG8000 1 w/v % | Dextran 2 w/v% | TEG 1 v/v % |
| Resolution range (Å) | 43.29-2.80 | 30.17-1.61 | 30.55-1.98 | 43.2-2.11 | 43.32-2.47 | 27.01-2.38 | 30.57-2.08 | 30.51-1.98 | 30.58-2.48 |
| Reflection used | 2,504 | 12,417 | 6,989 | 5,781 | 3,658 | 4,076 | 6,048 | 6,985 | 3,593 |
| R-factor (%) | 29.8 | 37.31 | 33.60 | 18.69 | 23.14 | 29.48 | 22.50 | 34.78 | 30.21 |
| Free R-factor (%) | 43.8 | 41.29 | 39.00 | 22.24 | 28.96 | 40.16 | 26.76 | 40.78 | 36.48 |
| R.m.s.deviations from ideal | | | | | | | | | |
| Bond length (Å) | 0.015 | 0.010 | 0.010 | 0.009 | 0.011 | 0.067 | 0.010 | 0.012 | 0.010 |
| Angle (°) | 1.99 | 1.50 | 1.40 | 1.16 | 1.64 | 3.25 | 1.29 | 1.38 | 1.45 |
| Ramachandran plot (%) | | | | | | | | | |
| Favored region | 70.73 | 94.12 | 91.18 | 97.75 | 88.10 | 94.37 | 96.55 | 94.03 | 98.33 |
| Allowed region | 19.51 | 5.88 | 8.82 | 2.25 | 9.52 | 5.63 | 3.25 | 5.97 | 1.67 |
| Outlier region | 9.76 | 0.00 | 0.00 | 0.00 | 2.38 | 0.00 | 0.00 | 0.00 | 0.00 |
